# Engineering *Y. lipolytica* for the biosynthesis of geraniol

**DOI:** 10.1101/2023.04.30.538875

**Authors:** Ayushi Agrawal, Zhiliang Yang, Mark Blenner

## Abstract

Geraniol is a monoterpene with wide applications in the food, cosmetics, and pharmaceutical industries. Microbial production has largely used model organisms lacking favorable properties for monoterpene production. In this work, we produced geraniol in metabolically engineered *Yarrowia lipolytica*. First, two plant-derived geraniol synthases (GES) from *Catharanthus roseus* (Cr) and *Valeriana officinalis* (Vo) were tested based on previous reports of activity. Both wild type and truncated mutants of GES (without signal peptide targeting chloroplast) were examined by co-expressing with MVA pathway enzymes tHMG1 and IDI1. Truncated CrGES (tCrGES) produced the most geraniol and thus was used for further experimentation. The initial strain was obtained by overexpression of the truncated HMG1, IDI and tCrGES. The acetyl-CoA precursor pool was enhanced by overexpressing mevalonate pathway genes such as ERG10, HMGS or MVK, PMK. The final strain overexpressing 3 copies of tCrGES and single copies of ERG10, HMGS, tHMG1, IDI produced approximately 1 g/L in shake-flask fermentation. This is the first demonstration of geraniol production in *Yarrowia lipolytica* and the highest de novo titer reported to date in yeast.

## 1. Introduction

Geraniol is an acyclic monoterpene alcohol obtained from aromatic plants and is widely used in fragrance and pharmaceutical industries (Chen and Viljoen, 2010). Geraniol exhibits anti-microbial (Lira et al., 2020), anti-fungal (Pattnaik et al., 1997) and insect-repellant properties (Barnard and Xue, 2004), and has been reported to have anti-cancer effects (Kim et al., 2012). Geraniol also serves as a precursor to strictosidine which is a central intermediate for over 3,000 monoterpene indole alkaloids (MIAs) (Brown et al., 2015). The titer in natural plants is low and the extraction process is laborious and expensive (Luo et al., 2015). Therefore, geraniol and other monoterpenoids are being synthesized in microbial cell factories like *E. coli* and *S. cerevisiae* using metabolic engineering strategies (Fischer et al., 2011; Liu et al., 2016). Yeasts are the preferred microbial cell factories for many natural products, due to the presence of eukaryotic organelles, the ability to produce organelle associated enzymes (e.g., P450s) (Wang et al., 2018) and the ability to perform post-translational modifications. Furthermore, yeast do not have natural phage, and tolerate low pH and a variety of environmental stresses (Raab and Lang, 2011) making them more robust in industrial practice. While *S. cerevisiae* is commonly used to demonstrate production of natural products (Y. Liu et al., 2019; Nielsen, 2014; Paddon et al., 2013; Palmer and Alper, 2019) due to the availability of synthetic biology and genetic engineering tools, it suffers from having low natural flux of natural product biosynthesis, the Crabtree effect that utilizes most available acetyl-CoA, limited availability of aromatic acids, and limited secretion systems (Nielsen, 2014; Palmer and Alper, 2019).

The oleaginous yeast *Yarrowia lipolytica* is a promising alternative to *S. cerevisiae* because it has a higher acetyl-CoA metabolic flux (Y. Liu et al., 2019), a much greater proportion of endogenous cytochrome P450 enzymes (Córdova et al., 2017), evolved to accumulate hydrophobic substrates (Fickers et al., 2005), lacks the Crabtree effect (Madzak, 2021), and has a growing array of metabolic engineering tools such as CRISPR-Cas9 (Ganesan et al., 2019; Gao et al., 2018; Schwartz et al., 2019, 2016), promoter libraries and, safe insertion of genes (Schwartz et al., 2017). These attributes combined position *Y. lipolytica* as a promising host for plant natural product synthesis. Indeed, numerous terpenoid products have been produced at high titers in *Y. lipolytica* and often higher titers with less extensive engineering compared to *S. cerevisiae* (e.g., α-farnesene, lycopene, β-carotene, astaxanthin, and α-humulene (Guo et al., 2022; Y. Liu et al., 2019; Ma et al., 2022; Zhu et al., 2022)).

Geraniol production has been demonstrated in *S. cerevisiae*. Early work focused on enhancing the mevalonate pathway flux, (Liu et al., 2013), followed by enhancing the supply of geranyl diphosphate selection of geraniol synthases and fusion to a geranyl pyrophosphate favoring mutant of the farnesyl-diphosphate synthase, resulting in modest titers (Zhao et al., 2016).Focusing on alternative sources of geraniol synthase, activating truncations of geraniol synthase, combined with ERG20 mutants favoring geranyl pyrophosphate, the highest shake flask titers in a eukaryote were reported (523.96 mg/L) (Jiang et al., 2017). Production in Yarrowia lipolytica has not be reported to date.

In this work, we engineered the mevalonate pathway to produce geraniol in biphasic cultures in metabolically engineered *Yarrowia lipolytica*. We screened two different types of geraniol synthases (GESs) from plants and determined truncated tCrGES performed best. The base strain was then engineered to increase the flux for geranyl pyrophosphate (GPP) synthesis by random integration of truncated HMG1, IDI1 and tCrGES. To enhance geraniol production, we further enhanced the precursor pool of acetyl-CoA by overexpressing the mevalonate pathway genes ERG10, HMGS, ERG12, ERG8 and optimized the truncated geraniol synthase gene copy number. Finally, process parameters such as carbon to nitrogen ratio in media were optimized for increasing the geraniol titers. However, optimizing the carbon to nitrogen ration by varying the nitrogen content did not significantly increase the geraniol titers. Instead, increasing the amount of glucose concentration led the titer to increase around 1 g/L in shake-flask cultivation.

## 2. Materials and methods

### 2.1 Plasmid construction

*E. coli* NEB 5α or *E. coli* NEB 10β was used for plasmid construction and propagation. Bacteria was grown in LB (Luria-Bertani) medium at 37 °C and 250 rpm with 100 mg/L ampicillin to select for all plasmids. Plasmids were assembled through SLIC or Gibson assembly. Shuttle vector pYLXP’ with pTEFintron promoter, XPR2 terminator, ampicillin marker for bacterial selection and Leu2/Ura3 markers for yeast selection was used as the starting vector which was kindly donated by Professor Peng Xu (UMBC, Maryland, USA). The genes of interest were cloned between the pTEFintron promoter and the XPR2 terminator to create a gene cassette. The heterologous genes *E. coli* AtoB, *C. roseus* GES, *V. officinalis* GES were codon-optimized for *Y. lipolytica* using the Genscript codon-optimization tool.

Constructs pYLXP’-EcAtoB, pYLXP’-ylHMGS, pYLXP’-ylHMG1, pYLXP’-ERG12 (MVK), pYLXP’-ERG8 (PMK), pYLXP’-ylIDI1, pYLXP’-ylGPPS, pYLXP’-CrGES, pYLXP’-VoGES from **SI Table-1** were cloned using Gibson assembly in pYLXP’ vector between pTEFintron promoter and XPR2 terminator. Signal peptide of gene GES (chloroplastic) was identified using Signal P 5.0/ Uniprot and with Jiang et al. work(Jiang et al., 2017) and truncated signal peptide version of the gene from two plant variants *C. roseus* (truncated 129 bp) and *V. officinalis* (truncated 138 bp) were also constructed in pYLXP’ vector by deleting the signal peptide using SLIC. 495 nucleotides from pYLXP’-HMG1 were truncated from N-terminal of HMG1 to generate t495HMG1 in pYLXP’ vector. Multi gene constructs (**SI Table-1**) pYLXP’-tCrGES-tHMG1-ylIDI, pYLXP’-EcAtoB-HMGS-tHMG1, pYLXP’-ylMVK-ylPMK, pYLXP’-EcAtoB-HMGS-tCrGES, pYLXP’-EcAtoB-HMGS-2xtCrGES were assembled using sub-cloning.

### 2.2 Strain construction

pYLXP’-tCrGES-tHMG1-ylIDI (Ura3) was linearized with NotI and transformed into *Y. lipolytica* PO1f cells to create strain S1 using the lithium acetate method as described in the next section. Post transformation, cells were plated on a YSC-Ura plate and incubated at 30 °C for 2 days. Multiple colonies were tested for geraniol production and the colony yielding the highest titer was selected for further engineering. Next, pYLXP’-EcAtoB-HMGS (Leu2), pYLXP’-ylMVK-ylPMK (Leu2), pYLXP’-EcAtoB-HMGS-tCrGES (Leu2) and pYLXP’-EcAtoB-HMGS-2xtCrGES (Leu2), pYLXP’-tCrGES-ylMVK-ylPMK (Leu2) were linearized with NotI and transformed into S1 cells using the lithium acetate method to create S2, S3, S4, S5 and S6 strains respectively, plated on YSC-Leu-Ura, and incubated at 30 °C for 2 days. Multiple colonies were tested for geraniol production and the colony yielding the highest titer was stored for further experimentation. The screened colonies were later grown in YPD media and genomic DNA was extracted using Thermo Fisher genomic DNA extraction kit. Integration was confirmed by PCR fragment size with gene specific primers (SI primer sequences) and subsequently by Sanger sequencing.

### 2.3 Medium and culture conditions

Geraniol synthase (GES) screening was performed on the plasmid bearing strains which were obtained by transforming the plasmids pYLXP’-tCrGES-tHMG1-IDI1, pYLXP’-CrGES-tHMG1-IDI1, pYLXP’-tVoGES-tHMG1-IDI1, pYLXP’-VoGES-tHMG1-IDI1 into *Y. lipolytica* PO1f using the lithium acetate method. In detail, PO1f was plated on solid YPD medium and incubated at 30 °C for 24 hours. Fresh one-step buffer for every individual reaction (100 µl) was prepared as follows: 90 µl of sterile 50% PEG 4000, 5 µl of sterile filtered 2 M lithium acetate, 2.5 µl of salmon sperm DNA (ssDNA was boiled at 98 °C for 7 minutes and snap-chilled on ice prior to mixing) and 2.5 µl of 2M dithiothreitol (DTT). Some amount of solid PO1f was picked up using an inoculation loop and mixed with the one-step buffer and vortexed for about 10 seconds. 300-500 ng of plasmid DNA was then added to the reaction and vortexed for 10 seconds. The tubes were then incubated at 39 °C for 1 hour. After 1 hour, 100 µl of the autoclaved MilliQ water was added to all the tubes and 100 µl mixture was plated on the selective plates and incubated at 30 °C for 2 days (this protocol is modified from Xu and Zhou lab protocol(Lv et al., 2019)). Fresh individual *Y. lipolytica* colonies were used for inoculation of pre-cultures in 14 ml culture tubes with 2 ml of selective medium with 40 g/L glucose. The pre-cultures were incubated at 28 °C, 220 rpm for 36-38 hours. 250 ml baffled flasks containing 50 ml of selective media (with respective dropouts) were inoculated with pre-cultures to a starting OD600 of 0.1. The culture was mixed well and 300-500 µl of the cultures was taken out and filtered using a 0.22 µm filter and run on HPLC-RID to check for initial glucose concentrations (**see Fig S3 and SI Table-3**). 10% overlay (5 ml isopropyl myristate; IPM) was added for in situ geraniol extraction and cultures were incubated at 28 °C, 220 rpm for 120 hours. After 120 hours, the overlay containing geraniol was analyzed by GC-MS (**see Fig S4 and Fig 2**) whereas the bottom layer was filtered and analyzed by HPLC-RID (**see SI Table-3**).

Multiple colonies were tested for strains S1 - S6. After *Y. lipolytica* transformation in the desired strain using the lithium acetate method, 8-10 colonies of each strain were picked and tested for geraniol production using the same fermentation method as described above. The colony with the highest geraniol titer was selected for further experimentation and to make a glycerol stock. The glycerol stocks of all strains were grown in triplicates to efficiently replicate the geraniol titers.

To optimize the media, the “one factor at a time” technique was used. Strain S6 was directly inoculated into 50 ml baffled flasks with 10 ml selective media as pre-cultures, followed by shake flask fermentation as described above. The C/N ratio was optimized by fixing glucose concentration (60 g/L) and varying ammonium sulfate concentrations (5 g/L, 8 g/L, 15 g/L, 30 g/L) to check the C/N ratios: 12:1, 7.5:1, 4:1, 2:1.

### 2.4 Biomass determination

The cell growth was monitored by measuring the optical density at 600 nm with an Eppendorf BioPhotometer (UV spectrophotometer). For dry cell weight (DCW, g/L) measurements, 5 ml of the well mixed cultures were transferred to 15 ml centrifuge tubes and centrifuged at 4,000 rpm for 5 minutes at room temperature and washed twice with sterile MilliQ water before re-suspending the pellet in 2 ml of sterile MilliQ water. The re-suspended pellets were transferred to pre-weighed aluminum trays and incubated in a vacuum oven until dry. The dried pellets in aluminum trays were weighed on a microbalance and the empty weight of the trays was subtracted.

### 2.5 Extraction of metabolite and quantification using analytical methods

After 120 hours of fermentation, the cultures were transferred to 50 ml centrifuge tubes and allowed to settle for 5-10 minutes at room temperature, after which two separate layers are observed. The top layer contained IPM with monoterpenoids and the bottom layer was aqueous media with cells. 1 ml of the IPM layer was collected and centrifuged at 15,000 rpm for 5 minutes at room temperature. After centrifugation, the top layer was transferred to GC vials. GC-MS (Thermo Fisher Trace 1300 GC and ISQ7000 mass spectrometer) equipped with TG-5MS column (30 m X 0.25 µm X 0.25 µm, Catalog number: 26098-1420) was used to detect monoterpenoids with helium as the carrier gas at a split ratio of 100:1 and 0.5 µl injection volume. The column temperature profile was 50 °C for 1 minute, 10 °C/min ramping to 250 °C and holding for 2 minutes. 1 mL of the aqueous (bottom) layer was collected and centrifuged at 15,000 rpm for 5 minutes at room temperature. The clarified aqueous solution was filtered using a 0.22 µm filter and run on HPLC-RID (Agilent 1290 Infinity II/Agilent 1100) for glucose measurements. For Agilent 1290, Agilent Hi-Plex H (PL1170-6830) was used along with guard column (PL1170-1830) for separation of glucose. The program was used as follows: 0.005 M sulfuric acid as mobile phase, flow rate of 0.4 ml/min, column compartment at 55 °C, RID at 55 °C, run time = 30 minutes and an injection volume of 10 µl. For Agilent 1100, Bio-Rad Aminex HPX 87H was used with 0.005 M sulfuric acid as mobile phase (Buday et al., 1990; Jones et al., 2016).

### 2.6 Quantitative PCR

GES gene copy number was evaluated using qPCR. The S6 strain was grown in YPD media and extracted during exponential phase using the Thermo Scientific yeast DNA extraction kit. Primer 3 (https://bioinfo.ut.ee/primer3-0.4.0/) was used for GES qPCR primer design (**see SI Figure-7**). The primer efficiency was between 90-110% (**see SI Figure-7**). 1 µl of genomic DNA was subjected to qPCR using the qScript™ One-Step SYBR® Green qRT-PCR Kit, QuantaBio from VWR (Catalog number: 95054-946) on CFX96 Real-Time PCR thermocycler. The following thermal profile was adopted: 50 °C for 10 min, 95 °C for 10 min, 95 °C for 15 s, 60 °C for 25 s, 72 °C for 30 s for 44 cycles. Single product amplification was verified by post amplification analysis of the melt curve. Cq value from the reaction was inserted into the standard curve equation. Copy number identified from the standard curve was divided by the number of genomes based on the mass of DNA loaded for the reaction and the size of the *Y. lipolytica* genome.

## 3. Results

The synthesis of monoterpenes requires geranyl pyrophosphate (GPP) produced by the condensation of isopentenyl pyrophosphate (IPP) and dimethylallyl pyrophosphate (DMAPP) in the mevalonate pathway (**Figure 1**). GPP is further condensed with IPP by ERG20 to form farnesyl diphosphate (FPP), the central precursor for sesquiterpenes and ergosterol, which is essential for cell growth (Zhao et al., 2016). Since ERG20 is essential to cell growth, it cannot be knocked out or it would result in cell death (Anderson et al., 1989). Hence, it is more challenging to produce monoterpenoids compared to sesquiterpenes pathway where the central intermediate is FPP. The toxicity of monoterpenes limits their production, with engineered yeast strains in many studies producing as little as 5 mg/L (Carter et al., 2003; Fischer et al., 2011). In order to alleviate the toxicity of monoterpenes to microorganisms, two-phase cultures are used with extraction solvents such as isopropyl myristate (Zeng et al., 2020), resulting in significantly higher titers (Brennan et al., 2012).

**Figure 1.**
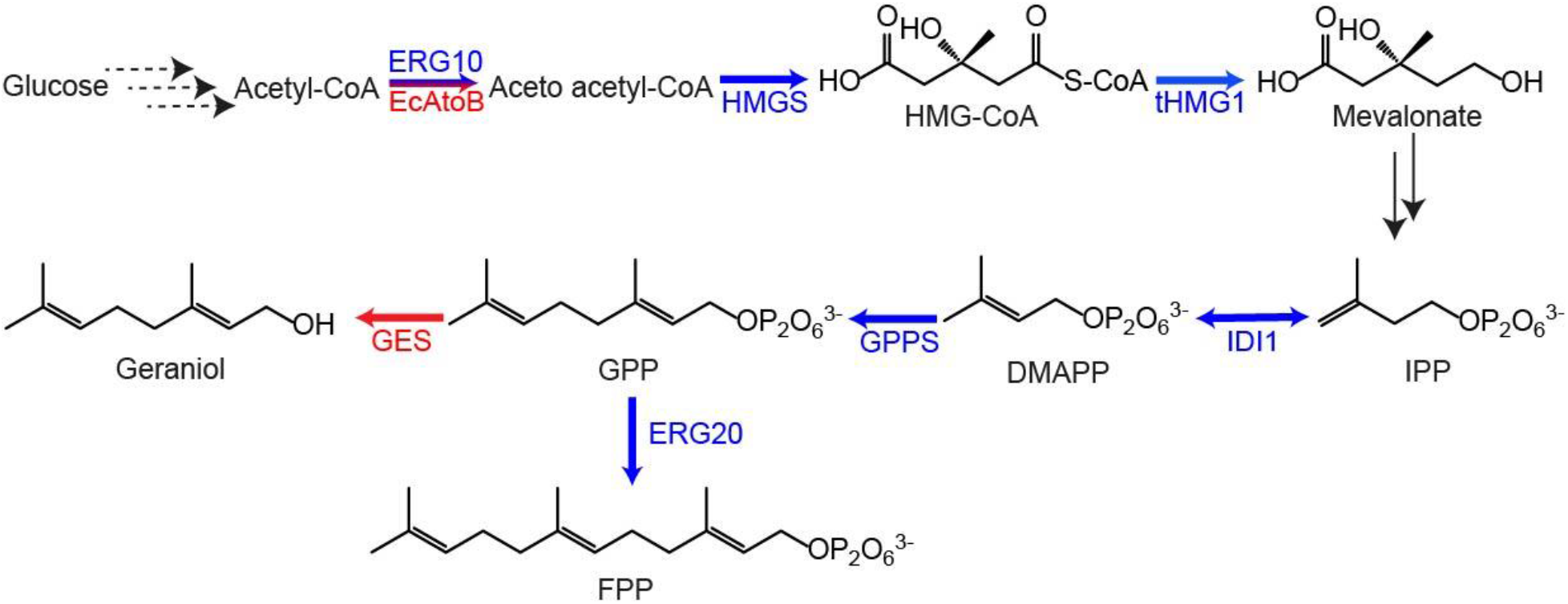
*Yarrowia lipolytica* geraniol biosynthesis pathway. The blue arrows indicate native enzymes and red arrows indicate heterologous enzymes. ERG10/AtoB is acetyl-CoA acetyltransferase, HMGS is HMG synthase, HMG is β-Hydroxy β-methylglutarate, tHMG1 is truncated HMG-CoA reductase, IPP is isopentyl 5-pyrophosphate, IDI1 is IPP isomerase, DMAPP is dimethylallyl 5-pyrophosphate, GPPS is geranyl pyrophosphate synthase, GES is geraniol synthase.

### 3.1 Testing of geraniol synthases (GESs)

To establish a geraniol biosynthetic pathway in *Y. lipolytica*, we first tested geraniol synthases from two plant sources. Geraniol synthase is natively localized in the chloroplast in plant cells; however, since yeast lacks chloroplasts, it is unclear where the enzyme localizes in the yeast cells. However, the localization can be confirmed using GFP tagging and then visualizing under a microscope. Jiang et al. screened geraniol synthases from nine plant sources for geraniol production in *S. cerevisiae* and identified *Catharanthus roseus* and *Valeriana officinalis* as the highest geraniol producing variants (Jiang et al., 2017). Jiang et al. further truncated the signal peptide of GES and found a several fold increase in geraniol titers compared to the non-truncated version in *S. cerevisiae* and discovered that the reason for higher titers is better protein folding as localization was not affected in truncated vs non-truncated forms as confirmed by RFP tagging. Hence, to test geraniol production in *Y. lipolytica*, we selected GES from plants *Catharanthus roseus* and *Valeriana officinialis* and tested truncated forms tCrGES (truncation ; strain S43) and tVoGES (strain S46), or expressed with native chloroplastic signal peptides, and co-expressed with tHMG1 and IDI1 as plasmids in PO1f cells. For non-truncated strains, CrGES and VoGES titers were similar (0.78± 0.2 mg/g of geraniol) whereas the truncation of the signal peptide increased the geraniol titers 6.4-fold (5± 1.3 mg geraniol/g DCW) for tCrGES and 4.6-fold (1.81± 0.52 mg/g of geraniol) for tVoGES compared to the non-truncated versions indicating that truncation is essential to higher geraniol titers. Among the truncated strains, tCrGES titer was 2.7-fold higher than tVoGES. Therefore, tCrGES was selected for further engineering of geraniol producing strains in *Y. lipolytica*.

### 3.2 Enhancement of MVA pathway genes and optimization of GES copy number

Since the MVA pathway is native to *Y. lipolytica* (G. Zhang et al., 2022), we tested whether the overexpression of MVA pathway genes (Arnesen and Borodina, 2022; Zhao et al., 2016) HMGS, HMG1, IDI1, MVK, PMK, and heterologous EcAtoB (which performs the same function as ERG10) would improve geraniol production by potentially increasing the GPP precursor pool. Overexpression of EcAtoB was pursued as it has better activity than the endogenous ERG10 (Y. Liu et al., 2019; Martin V et al., 2003). In the MVA pathway, HMG1 gene that catalyzes the conversion of HMG-CoA to mevalonate is identified as a metabolic bottleneck due to its limited activity (Donald et al., 1997). Hence, the endoplasmic reticulum (ER) targeting signal peptide of HMG1 (first 495 nucleotides at the N-terminal) were truncated to improve the enzyme activity (Liu et al., 2020). Moreover, extra copies of tCrGES were overexpressed to divert the flux to geraniol synthesis.

From GES testing, plasmid containing tCrGES, tHMG1 and IDI1 was linearized and transformed to PO1f strain. The resulting strain S1 produced 119±7.85 mg geraniol/L or 22±1.74 mg geraniol/g DCW (**Figure-3**), possibly due to stable expression due to gene integration compared to being expressed as plasmids as in **Figure-2**. Strain S1 was selected as the base strain to construct the rest of the strains for this work. Strain S2 was constructed by linearizing the plasmid containing EcAtoB and HMGS genes and transforming to S1 strain. Strain S2 produced 277.73±11.3 mg geraniol/L or 40.6±3.4 mg geraniol/g DCW. EcAtoB and HMGS genes were potentially able to enhance the formation of geraniol by increasing production of the direct precursor GPP. The flux of GPP pushed towards geraniol production led to an increase in geraniol titer by 1.8-fold mg geraniol/g DCW. Strain S3 was constructed by linearizing the plasmid containing ERG12 (MVK) and ERG8 (PMK) genes and transforming into S1 strain. Strain S3 produced 207.56±14 mg geraniol/L or 27.5±7.5 mg geraniol/g DCW.

**Figure 2.**
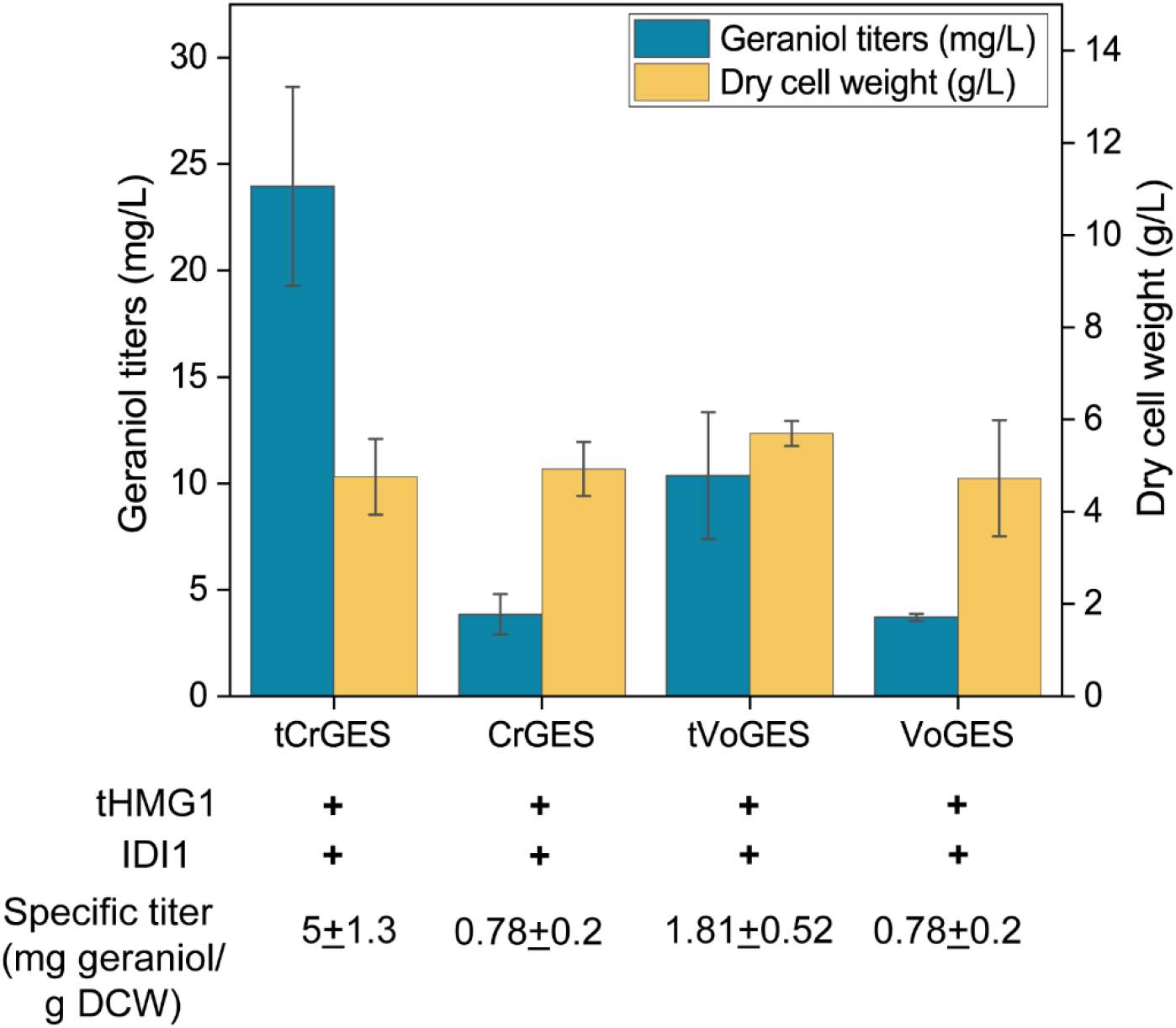
Truncation of chloroplast signal peptide leads to higher geraniol titers. Geraniol titers (cyan) were significantly higher after truncation of the chloroplast signal peptide (t). The dry cell weights (yellow) were not affected by GES expression (**see SI Table-3**). GES: Geraniol synthase, Cr: *Catharanthus roseus*, Vo: *Valeriana officinalis*, tHMG1 is truncated HMG reductase, IDI1 is *Yarrowia lipolytica* IPP isomerase. Error bars are standard deviation of three biological triplicates.

**Figure-3:**
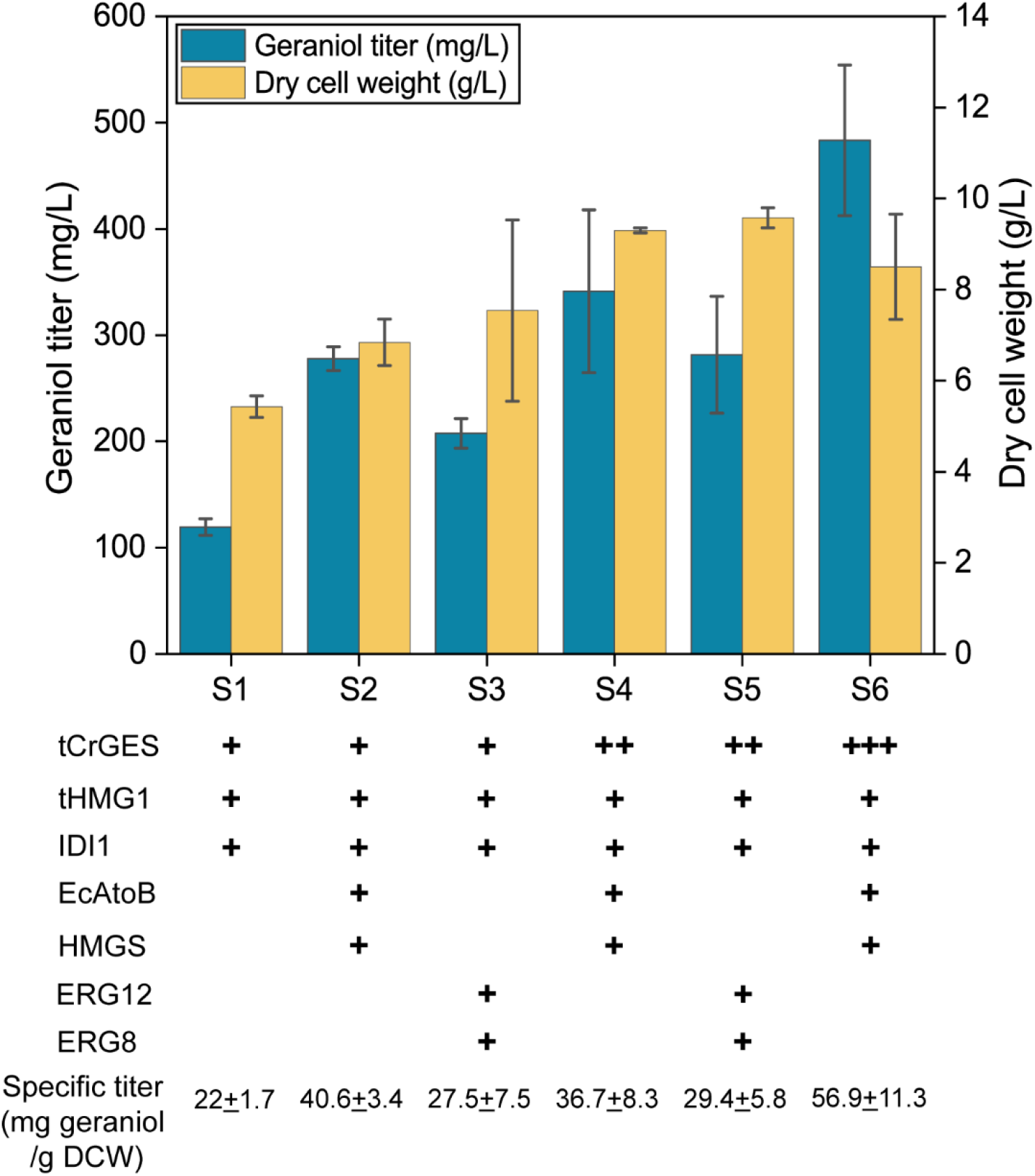
Strain engineering for geraniol production in *Yarrowia lipolytica*. The base strain was obtained by overexpression of the tHMG1, IDI and tCrGES. The precursor pool of acetyl-CoA was enhanced by overexpression of the MVA pathway genes such as ERG10, HMGS or ERG12, ERG8. The effect of tCrGES copy number was also investigated. The final strain overexpressing three copies of tCrGES and single copies of ERG10, HMGS, tHMG1, IDI can produce 483 mg/L of geraniol in shake flask cultivation. Error bars are standard deviation of three biological replicates.

The specific titer of geraniol produced in strain S3 reduced by 1.5-fold compared to S2 but was still 1.25-fold higher when compared to S1 strain. Although, the overexpression of combinations of genes EcAtoB/HMGS and MVK/PMK both increased geraniol titers, the overexpression of EcAtoB/HMGS turned out to be better for higher geraniol production than MVK/PMK. Strain S4 was constructed by linearizing the plasmid containing tCrGES, EcAtoB and HMGS and transforming to S1 strain. Strain S4 produced 341.17±76.7 mg geraniol/L or 36.7±8.2 mg geraniol/g DCW. Strain S5 was constructed by linearizing the plasmid containing tCrGES, ERG12 and ERG8 and transforming into Strain S1. Strain S5 produced 281.37±54.91 mg geraniol/L or 29.4±5.8 mg geraniol/g DCW. The titer of geraniol increased when compared to S1 but was not significantly different from S4 as observed with strain S3 as well concluding the overexpression of EcAtoB/HMGS is more helpful in increasing geraniol titers when compared to the overexpression of MVK/PMK genes. Hence, we engineered strain S6 which was obtained by linearizing the plasmid containing 2 copies of tCrGES, and one copy of each EcAtoB and HMGS and transforming to S1 strain. The gene combination was selected because the titers increased significantly when comparing S2 and S1 strains. Also, the extra tCrGES could help push the flux through the geraniol pathway. As expected, the titers of geraniol were higher compared to S1, S2, S3, S4 or S5 strains. Strain S6 produced 483.39±70.8 mg geraniol/L or 56.9±11.3 mg geraniol/g DCW which was 1.55-fold mg geraniol/g DCW higher than strain S4 and 1.93 mg geraniol/g DCW higher than strain S5. Hence, strain S6 was selected for further optimization of the geraniol biosynthetic pathway.

### 3.3 Process parameters optimization

We observed that strains S4, S5 and S6 almost completely consumed the glucose (**see SI Table-3**) supplied at the start of the reaction (40 g/L). This implies that we may benefit from adding more glucose at the start of the reaction or by using a fed-batch strategy where we could supply some glucose after 2-3 days. Hence, we increased the initial glucose concentration to 60 g/L and simultaneously also varied the nitrogen content in media for media optimization.

To optimize the fermentation medium, one-factor at a time optimization technique was used. Carbon/nitrogen (C/N) ratio was optimized by keeping glucose constant at 60 g/L while varying the amount of ammonium sulfate concentrations (nitrogen concentrations) in the fermentation medium. C/N ratio is critical for regulating acetyl-CoA flux in *Y. lipolytica*(H. Liu et al., 2019). Hence, various C/N ratios 12:1, 7.5:1. 4:1 and 2:1 were tested. C/N ratio of 12:1, 7.5:1 and 4:1 all resulted in approximately 1 g/L geraniol with similar amounts of geraniol/DCW (mg/g) as shown in **Figure-4**. However, the amount of geraniol produced/DCW (mg/g) was significantly lower for ammonium sulfate concentration of 30 g/L. This could be because the excess nitrogen supplied may have diverted more carbon to cell growth which is consistent with a previous study for squalene production(Liu et al., 2020). Also, compared to the non-optimized medium for geraniol production in **Figure-3**, the titer was 3.2-fold higher (mg/g DCW) for the optimized media. Overall, the higher geraniol titer after process optimization could be attributed to the higher initial glucose concentration. Media optimization did not result in any significant increase in the geraniol titers.

**Figure-4:**
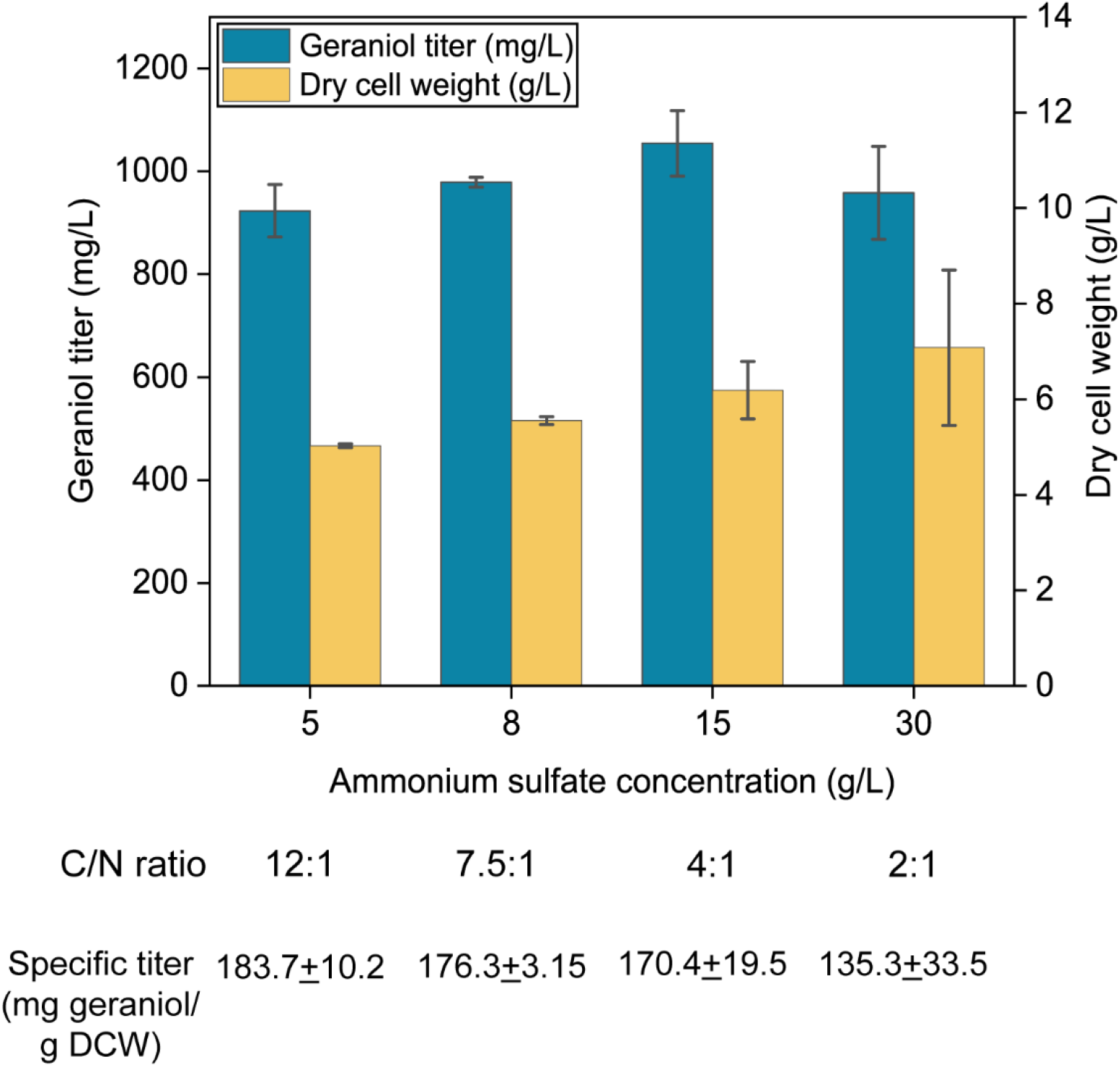
Process parameters optimization for geraniol production in *Yarrowia lipolytica*. Strain S6 was subjected to different C/N ratios by varying nitrogen content in the media in shake flask cultivation. All the ratios tested yielded around 1 g/L geraniol. Error bars are standard deviation of three biological replicates.

## 4. Discussion

In this work, *Yarrowia lipolytica* was engineered as a heterologous host for geraniol production. We first identified two geraniol synthases from plants *Catharanthus roseus* and *Valeriana officinalis* and confirmed prior findings that truncation of geraniol synthase was essential to increase geraniol titers (Jiang et al., 2017). Moreover, overexpression of MVA pathway genes and increasing tCrGES copy number increased the titers to 483 mg geraniol/L. Finally, the titer was increased to 1 g/L geraniol by increasing the glucose concentration in the media. To the best of our knowledge, this work represents the first time that geraniol has been produced in *Y. lipolytica*.

Many studies of geraniol production in *E. coli* and *S. cerevisiae* have been published based on overexpression of rate-limiting enzymes in the MVA pathway (Liu et al., 2013), dynamic control of ERG20 expression and OYE2 deletion (Zhao et al., 2017), increasing GPP pool, minimizing dehydrogenation of geraniol (Zhou et al., 2014) construction of fusion proteins to facilitate substrate channeling (Zhao et al., 2016), compartmentalization of the geraniol pathway in mitochondria (Yee et al., 2019), peroxisomal localization and improvement of geraniol tolerance (Gerke et al., 2020), optimization of fermentation conditions, and interconversion of geranyl acetate and geraniol (Liu et al., 2016) with titers less than 525 mg/L in shake-flask fermentation in most papers. Our shake-flask results of 1g/L geraniol exceeds shake-flask fermentation titers produced by these engineered *S. cerevisiae*, suggesting that *Y. lipolytica* is a preferred host for the production of monoterpenoids.

Recently, Zhao et al. engineered the non-conventional yeast *Candida glycerinogens* using dual pathway engineering with MVA pathway and the isopentenol utilization pathway, and by dynamic regulation using an inducible promoter (Zhao et al., 2022). Shake-flask titers using solely glucose remained under 500 mg/L. Therefore, our strain performs better when doing de novo geraniol biosynthesis from just glucose. However, they were able to achieve 1.2 g/L geraniol in shake-flask fermentation using a combination of glucose and prenol, and a decane inducible promoter. A focused effort to combine important advances, such as dynamic regulation, with the inherent beneficial properties of *Y. lipolytica* could further improve the high titers we achieved in this study.

Monoterpenoids are often toxic to microorganisms (Brennan et al., 2012). In this study, we found that around 200 mg geraniol/L completely inhibited the cell growth (**see Fig S1**). Hence, we used the biphasic fermentation system for geraniol production by adding 10% v/v isopropyl myristate to the cultures.

However, the addition of the overlay does not stop the conversion of geraniol to other monoterpenoids as is commonly observed (Zhao et al., 2017). We found that non-geraniol monoterpenoids (to linalool, nerol, citronellol and geranic acid) were formed with somewhat consistent titers by enzymatic and non-enzymatic conversions during the fermentation period (**see Fig S8**). The best producing strain under optimized conditions (**see SI-Table 2**) had 78-84% of monoterpenoids as geraniol rather than derivatives. In *S. cerevisiae*, OYE2 (Steyer et al., 2013) enzyme is the major contributor to the reaction of geraniol to citronellol and alcohol dehydrogenases are responsible for the conversion of geraniol to geranic acid (Ohashi et al., 2021). *Y. lipolytica* also natively harbors the old yellow enzymes (as confirmed by NCBI BLAST using *S. cerevisiae* OYE2 as the reference sequence) and alcohol dehydrogenases likely responsible for the enzymatic conversion of geraniol to citronellol and geranic acid (Gatter et al., 2014). Therefore, to further increase geraniol production, the enzymes responsible for enzymatic conversion could be knocked out. Non-enzymatic conversion has less obvious solutions; however, increasing flux to a downstream product may prevent its conversion to undesired products.

Geraniol is the first committed step towards monoterpene indole alkaloid (MIA) biosynthesis. Seminal work in engineering MIA focused on strictosidine, the first intermediate of the broad MIA class of compounds (Brown et al., 2015). Other papers have also demonstrated production of monoterpenes and strictosidine with key improvements achieved by pathway compartmentalization, using geraniol and tryptamine feeds, and optimizing P450 and P450 reductase coupling (Misa et al., 2022; Yee et al., 2019) *S. cerevisiae* was recently engineered to produce vincristine (Gao et al., 2022) and vinblastine (J. Zhang et al., 2022). Monoterpene indole alkaloid titers could be enhanced further by eliminating geraniol supply limitations through the combined approaches described in this paper and the use of a more favorable host such as *Y. lipolytica*.

## 5. Conclusions

In summary, we engineered a high geraniol producing strain in *Yarrowia lipolytica*, the first demonstration in *Y. lipolytica*. We have achieved the highest titers for de novo geraniol production from yeast, over 1 g/L. The work suggests a superior role for Y. lipolytica as a host for making monoterpenes and other derivatives. This strain can be engineered further through various metabolic engineering strategies to produce more geraniol or used as a starting point for monoterpene indole alkaloids with therapeutic applications.

## 6. Author contributions

**Ayushi Agrawal**: Conceptualization, Formal analysis, Investigation, Methodology, Validation, Visualization, Writing – original draft. **Zhiliang Yang**: Conceptualization, Formal analysis, Investigation, Methodology, Writing – review & editing. **Mark Blenner**: Conceptualization, Funding acquisition, Project administration, Supervision, Writing – review & editing.

## 8. Acknowledgements

The authors would like to acknowledge Phil O’Dell, Vijay Ganesan, and Anthony Stohr for insightful comments on the manuscript. The authors would also like to knowledge E. Terry Papoutsakis for access to his group’s HPLC at times throughout the project. Research reported in this publication was supported by the National Institute of General Medical Sciences of the National Institutes of Health under award number 1R35GM133803.

