## Supplementary material for "Engineering *Y. lipolytica* for the biosynthesis of geraniol": SI Table-3

**Fig S1. Geraniol toxicity assessment**

From previous reports of geraniol production in *S. cerevisiae*, it was found that the cell growth was significantly hindered for geraniol concentration greater than 200 mg/L. Hence, we performed the geraniol toxicity assessment experiment to evaluate the toxicity of geraniol in *Y. lipolytica* cells. S6 (3xtCrGES-tHMG1-ylIDI1-EcAtoB-ylHMGS) strain cells were streaked on a YPD plate and grown for 24 hours at 30 °C. A single colony from the plate was inoculated in YPD media for 24 hours at 30 °C. The OD600 of the cells was measured and the cells were washed thrice with autoclaved MilliQ water. The OD600 of the cells was adjusted to 1 and the cells were serially diluted to $10^{-1}$, $10^{-2}$, $10^{-3}$, $10^{-4}$. 50 µl of the samples were plated on YPD plates containing 0, 100, 200, 350, 500 mg/L geraniol and the plates were incubated at 30 °C for 2 days. Around 200 mg/L geraniol inhibited the growth of the S6 strain in *Y. lipolytica* which is in agreement with the results observed in *S. cerevisiae*[1].

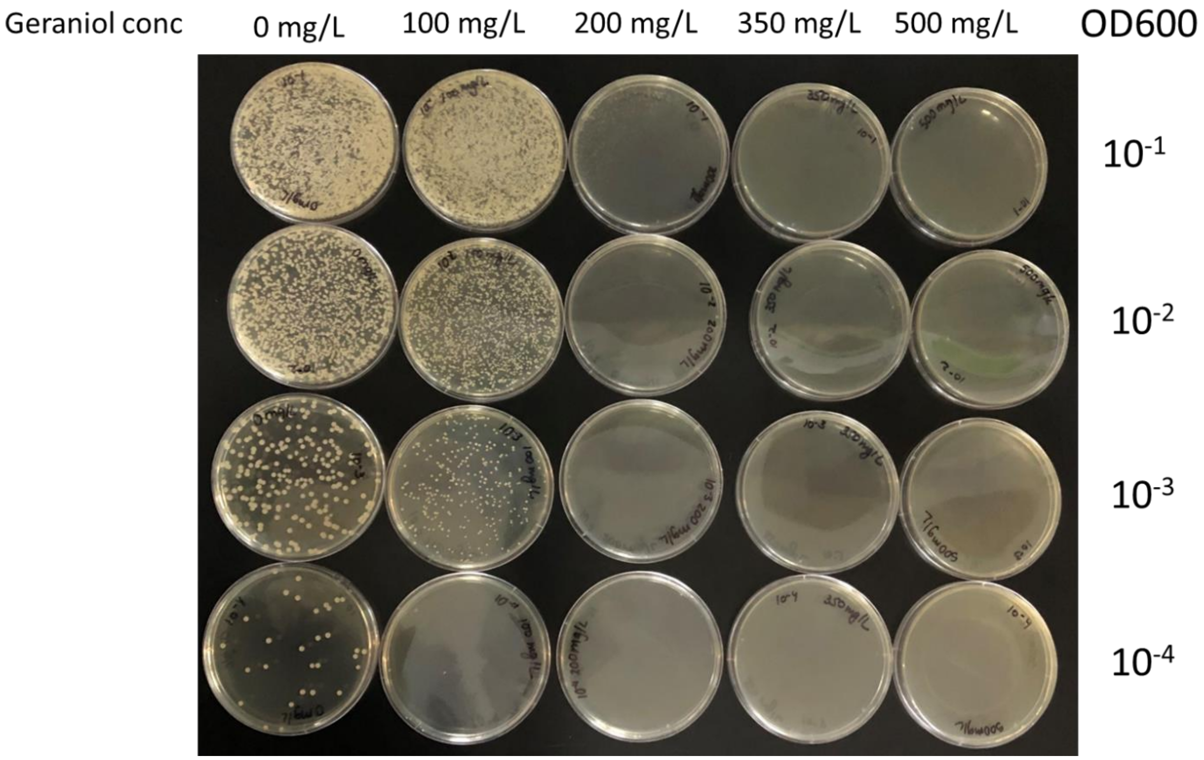

**Fig S2. Geraniol standard curve:** Fresh geraniol standards in isopropyl myristate were run in triplicates (three injections from the same vial) for every experiment.

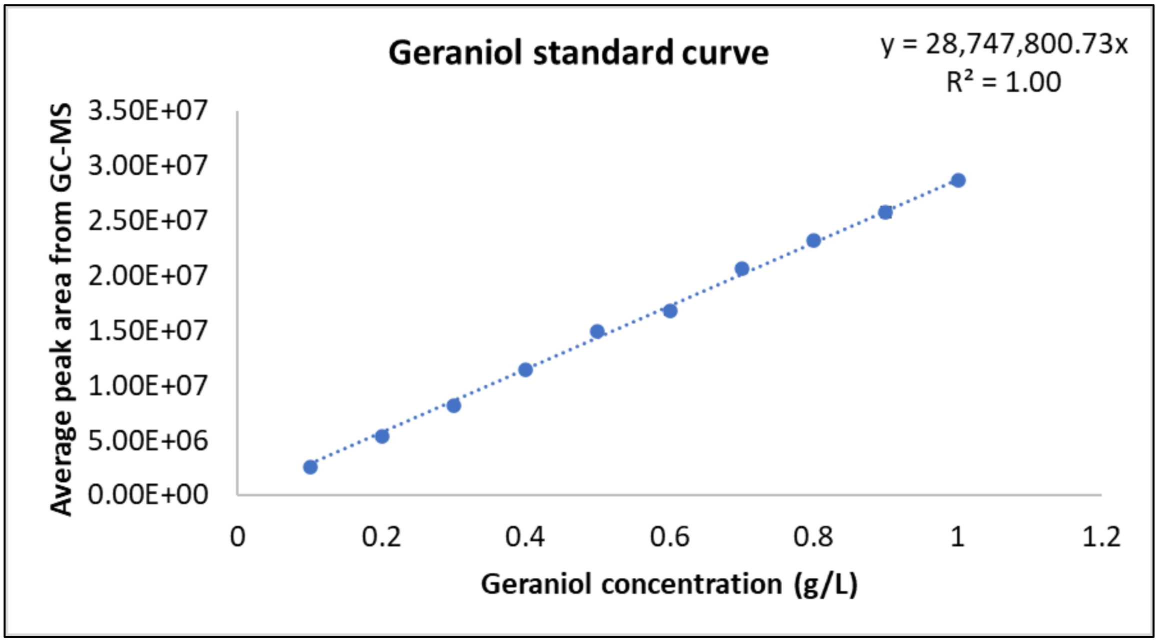

**Fig S3. Glucose standard curve:** Fresh glucose standards in water were run in triplicates (three injections from the same vial) for every experiment.

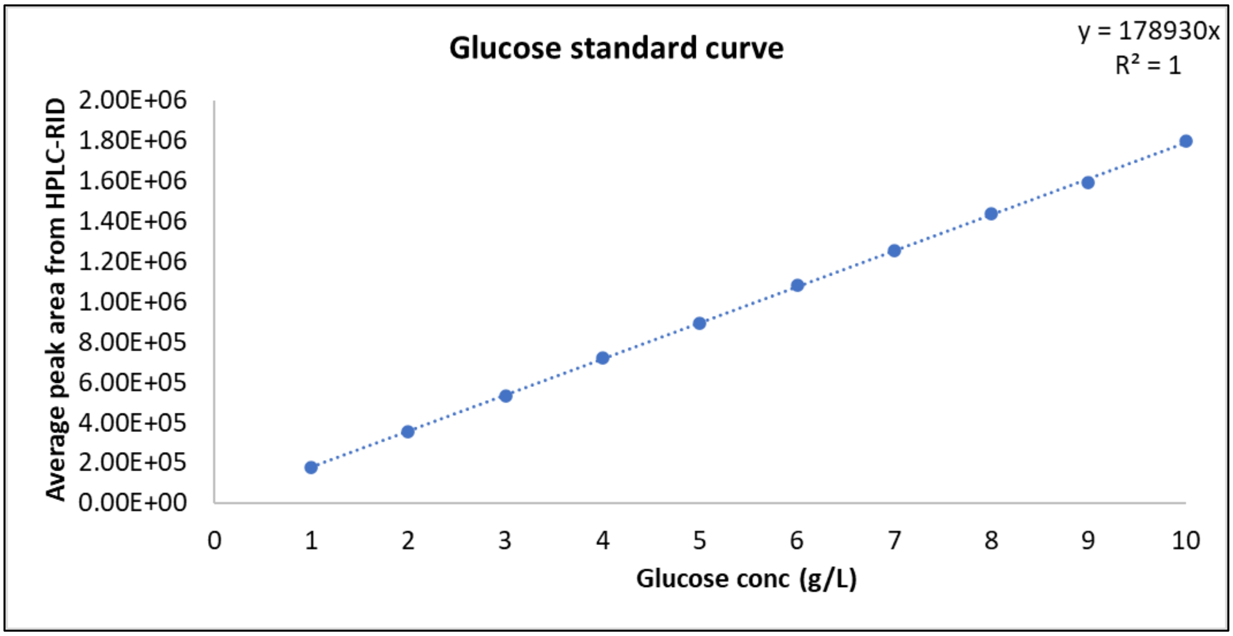

**Fig S4. Chromatogram of 500 mg/L geraniol standard dissolved in isopropyl myristate in GC-MS**

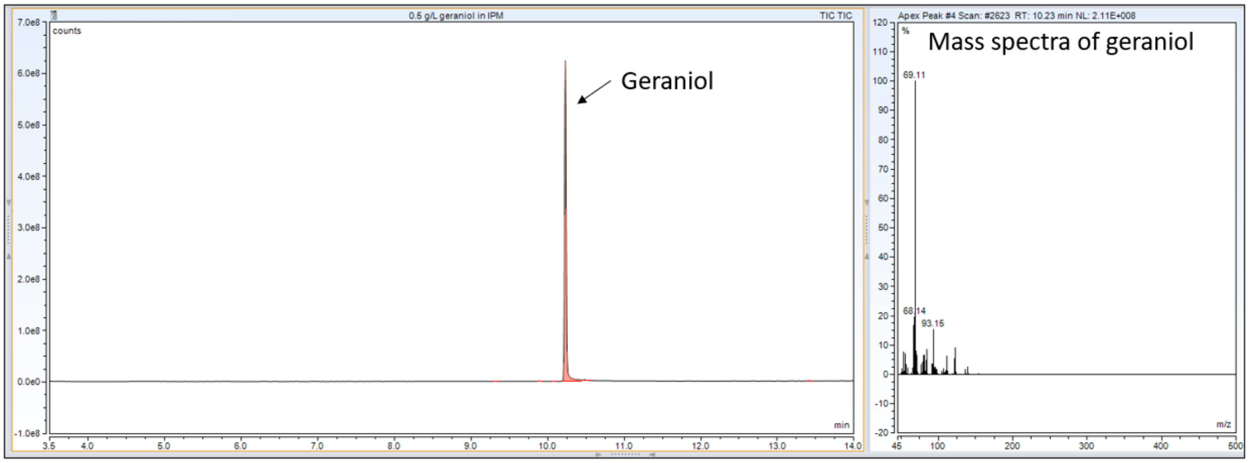

**Fig S5. Chromatogram of all monoterpenoids as identified by GC-MS in one of the samples**

For all monoterpenoids linalool, citronellol, nerol, and citral standards were purchased, and the identities were re-confirmed with GC-MS.

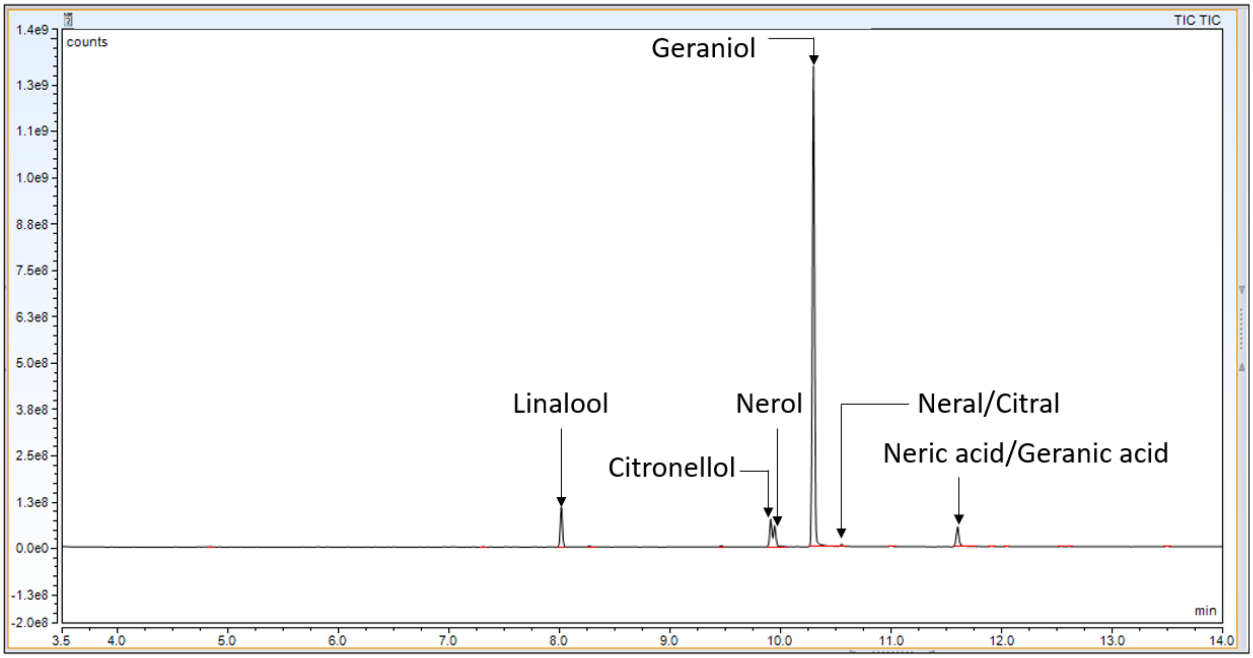

**Fig S6. Chromatogram of 2 g/L glucose from HPLC-RID using Agilent Hi Plex H column**

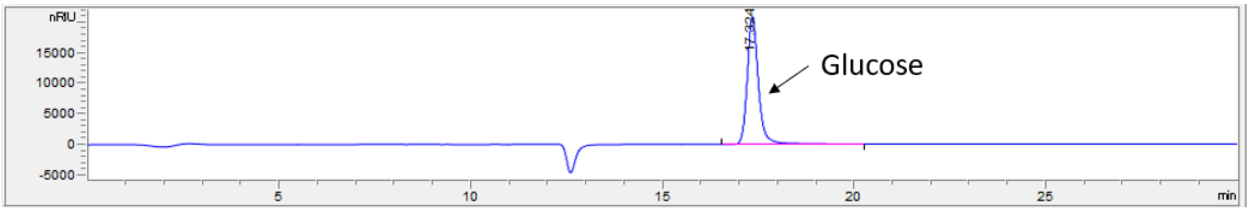

**Fig S7. Copy number analysis of the final strain S6**

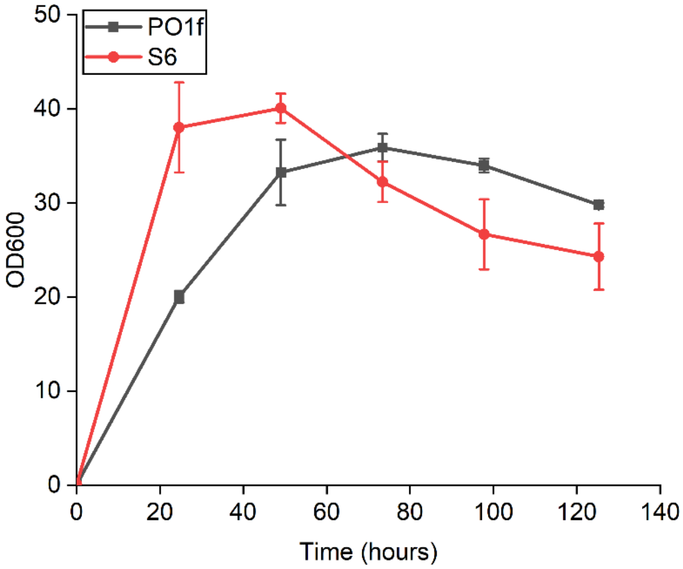

The final strain S6 and the *Yarrowia lipolytica* WT PO1f strain (control) were grown for a duration of over 120 hours to check for various growth phases. At the exponential phase, genomic DNA was extracted for the S6 strain and qPCR was performed to evaluate the tCrGES copy number.

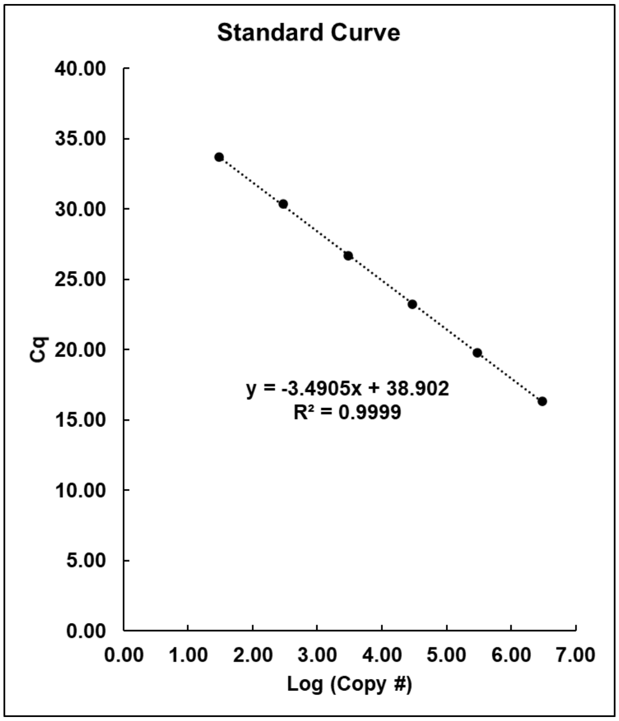

qPCR primers used: FP: CACCACCAACGAGATTTGCTAC, RP: TTGAACCACTCAGCCTCCAC

The primer efficiency was found to be 93.42%. Using this standard curve, the copy number of S6 strain was found to be 2.6±0.22. The copy number was evaluated as a result of 6 technical replicates.

**Fig S8. Geraniol and its conversion to other monoterpenoids**[2]

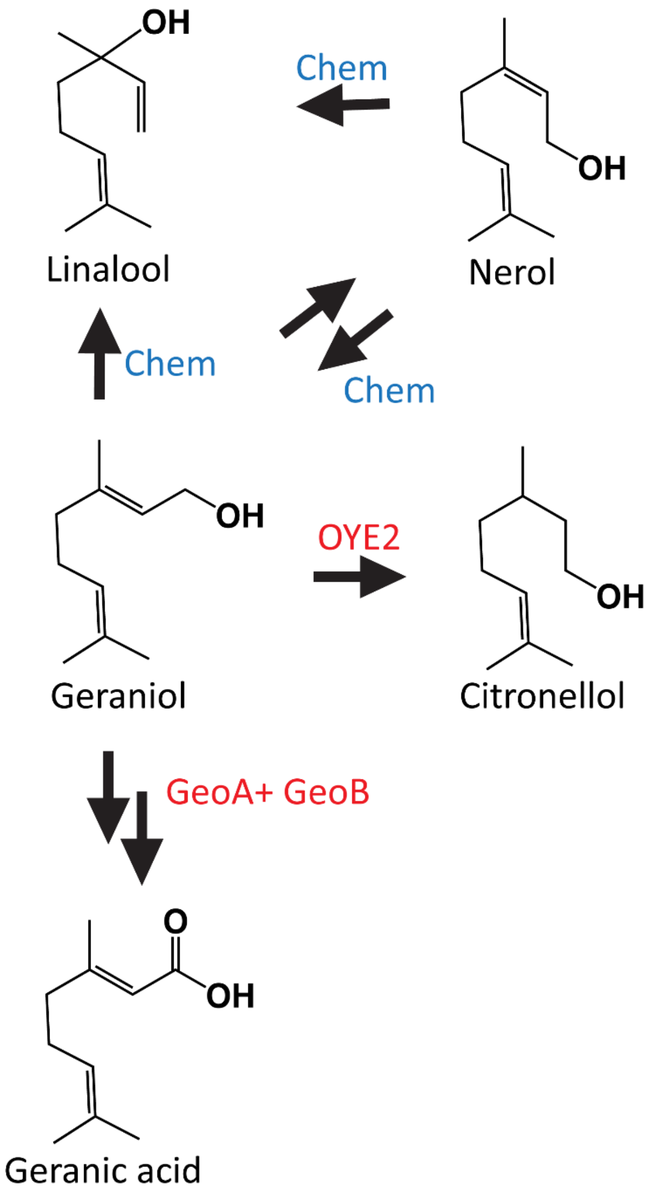

**OYE2:** old yellow enzyme, **GeoA**: geraniol dehydrogenase, **GeoB**: geranial dehydrogenase, Chem indicates chemical conversion.

**Table-1: Plasmids and strain list:**

| Plasmids | Genotype | Source |
|---|---|---|
| pYLXP’ | pYaliA1 vector backbone with leucine marker and Ampicillin resistance gene | Xu Peng et al |
| pYLXP'-EcAtoB | pYLXP' carrying AtoB from *E. coli* | This work |
| pYLXP'-ylHMGS | pYLXP' carrying HMGS from *Y. lipolytica* | This work |
| pYLXP'-yltHMG1 | pYLXP' carrying truncated HMG1 from *Y. lipolytica* | This work |
| pYLXP'-ERG12 (MVK) | pYLXP' carrying ERG12 from *Y. lipolytica* | This work |
| pYLXP'-ERG8 (PMK) | pYLXP' carrying ERG8 from *Y. lipolytica* | This work |
| pYLXP'-ylIDI1 | pYLXP' carrying IDI from *Y. lipolytica* | This work |
| pYLXP'-ylGPPS | pYLXP' carrying GPPS from *Y. lipolytica* | This work |
| pYLXP'-CrGES | pYLXP' carrying GES from *Catharanthus roseus* | This work |
| pYLXP'-tCrGES | pYLXP' carrying truncated GES from *Catharanthus roseus* | This work |
| pYLXP'-VoGES | pYLXP' carrying GES from *Valeriana officinalis* | This work |
| pYLXP'-tVoGES | pYLXP' carrying truncated GES from *Valeriana officinalis* | This work |
| pYLXP'-tCrGES-tHMG1-ylIDI (Ura3) | pYLXP' carrying truncated GES from *Catharanthus roseus*, truncated HMG1 from *Y. lipolytica* and IDI from *Y. lipolytica* | This work |
| pYLXP'-EcAtoB-HMGS (Leu2) | pYLXP' carrying AtoB from *E. coli* and truncated HMG1 from *Y. lipolytica* | This work |
| pYLXP'-ylMVK-ylPMK (Leu2) | pYLXP' carrying MVK and PMK from *Y. lipolytica* | This work |
| pYLXP'-EcAtoB-HMGS-tCrGES (Leu2) | pYLXP' carrying AtoB from *E. coli*, HMGS from *Y. lipolytica* and truncated GES from *Catharanthus roseus* | This work |
| pYLXP'-EcAtoB-HMGS-2xtCrGES (Leu2) | pYLXP' carrying AtoB from *E. coli*, HMGS from *Y. lipolytica* and 2 copies of truncated GES from *Catharanthus roseus* | This work |
| pYLXP'-tCrGES-ylMVK-ylPMK (Leu2) | pYLXP' carrying truncated GES from *Catharanthus roseus*, MVK and PMK from *Y. lipolytica* | This work |

| Strains | Genotype | Source |
|---|---|---|
| *E. coli* NEB 5α | fhuA2 Δ(argF-lacZ)U169 phoA glnV44 Φ80 Δ(lacZ)M15 gyrA96 recA1 relA1 endA1 thi-1 hsdR17 | New England Biolabs |
| *E. coli* NEB 10β | *Δ(ara-leu) 7697 araD139 fhuA ΔlacX74 galK16 galE15 e14- φ80dlacZΔM15 recA1 relA1 endA1 nupG rpsL* ($Str^R$) *rph spoT1 Δ(mrr-hsdRMS-mcrBC)* | New England Biolabs |
| *Y. lipolytica* PO1f | MatA, leu2−270, ura3−302, xpr2−322, axp-2 (ATCC no. MYA-2613) | Madzak C et al |
| S1 | PO1f cells harboring NotI digested pYLXP'-tCrGES-tHMG1-ylIDI | This work |
| S2 | PO1f cells harboring NotI digested pYLXP'-tCrGES-tHMG1-ylIDI and NotI digested pYLXP'-EcAtoB-HMGS | This work |
| S3 | PO1f cells harboring NotI digested pYLXP'-tCrGES-tHMG1-ylIDI and NotI digested pYLXP'-ylMVK-ylPMK | This work |
| S4 | PO1f cells harboring NotI digested pYLXP'-tCrGES-tHMG1-ylIDI and NotI digested pYLXP'-EcAtoB-HMGS-tCrGES | This work |
| S5 | PO1f cells harboring NotI digested pYLXP'-tCrGES-tHMG1-ylIDI and NotI digested pYLXP'-tCrGES-ylMVK-ylPMK | This work |
| S6 | PO1f cells harboring NotI digested pYLXP'-tCrGES-tHMG1-ylIDI and NotI digested pYLXP'-EcAtoB-HMGS-2xtCrGES | This work |

**Table-2: Concentration of total and other monoterpenoids other than geraniol in this work**

| Total monoterpenoid titers/DCW and Other monoterpenoid titers/DCW (mg/g) | | | | | |
|---|---|---|---|---|---|
| | **Plasmid** | **Total monoterpenoid concentration (mg/g DCW)** | **Std. Dev. (total)** | **Other monoterpenoid concentration (mg/g DCW)** | **Std. Dev. (other)** |
| **Figure-2** | pYLXP'-tCrGES-tHMG1-ylIDI (1) | 13.36151654 | 2.759545651 | 8.329153152 | 1.842090181 |
| | pYLXP'-CrGES-tHMG1-ylIDI (2) | 3.99959642 | 0.647451806 | 3.218764096 | 0.549907901 |
| | pYLXP'-tVoGES-tHMG1-ylIDI (3) | 3.435834103 | 0.183030962 | 2.785682774 | 0.153376684 |
| | pYLXP'-VoGES-tHMG1-ylIDI (4) | 9.273513473 | 2.708987851 | 7.082535323 | 2.095022652 |
| | **Strain** | **Total monoterpenoid concentration (mg/g DCW)** | **Std. Dev. (total)** | **Other monoterpenoid concentration (mg/g DCW)** | **Std. Dev. (other)** |
| **Figure-3** | S1 | 32.73011824 | 2.295085062 | 10.72055029 | 1.141879056 |
| | S2 | 91.21189892 | 9.318687697 | 50.60781336 | 7.22555211 |
| | S3 | 47.17521654 | 14.92012864 | 19.64700976 | 9.523699477 |
| | S4 | 76.59362965 | 11.72318284 | 39.90864369 | 8.317470988 |
| | S5 | 41.29937248 | 5.850037124 | 11.90766082 | 0.692972865 |
| | S6 | 90.16503921 | 19.06794982 | 33.2954753 | 12.80738901 |
| | **Ammonium sulfate concentration** | **Total monoterpenoid concentration (mg/g DCW)** | **Std. Dev. (total)** | **Other monoterpenoid concentration (mg/g DCW)** | **Std. Dev. (other)** |
| **Figure-4** | 5 g/L | 220.0683533 | 10.375734 | 36.38354206 | 1.445367483 |
| | 8 g/L | 226.6595809 | 5.391006451 | 50.39777142 | 3.889275543 |
| | 15 g/L | 213.0911808 | 23.26461649 | 42.65855183 | 4.988229544 |
| | 30 g/L | 169.301306 | 40.93856574 | 33.97284034 | 8.134960557 |

**Other monoterpenoids** include Linalool, Citronellol, Nerol, Neral/Citral, Neric acid/Geranic acid

**Total monoterpenoids** include Linalool, Citronellol, Nerol, **Geraniol**, Neral/Citral, Neric acid/Geranic acid

**Table-3: Glucose consumption chart for all figures as evaluated by HPLC-RID**

| Initial and final concentrations of glucose (g/L) | | | | | |
|---|---|---|---|---|---|
| Figure-2 | Plasmid | Glucose conc at 0 hours (g/L) | Std. Dev. (0 hours) | Glucose conc at 120 hours (g/L) | Std. Dev. (120 hours) |
| | pYLXP'-tCrGES-tHMG1-ylIDI | 40.89723967 | 2.06519967 | 15.96956948 | 2.56428921 |
| | pYLXP'-CrGES-tHMG1-ylIDI | 39.45821723 | 0.102992934 | 16.35992153 | 2.916722198 |
| | pYLXP'-tVoGES-tHMG1-ylIDI | 39.40627177 | 0.276129151 | 13.21761052 | 0.92039401 |
| | pYLXP'-VoGES-tHMG1-ylIDI | 39.47497949 | 0.20397545 | 17.45238109 | 1.878735384 |
| Figure-3 | Strain | Glucose conc at 0 hours (g/L) | Std. Dev. (0 hours) | Glucose conc at 120 hours (g/L) | Std. Dev. (120 hours) |
| | S1 | 40.0368346 | 0.696001078 | 13.3904392 | 0.280494592 |
| | S2 | 39.23798848 | 0.058253654 | 3.239113926 | 0.744016841 |
| | S3 | 40.15793955 | 0.300126819 | 4.478888555 | 3.850973763 |
| | S4 | 40.08523617 | 0.304902321 | 0 | 0 |
| | S5 | 43.4706226 | 0.361946303 | 1.3982446 | 0.877327023 |
| | S6 | 43.17333734 | 1.610957808 | 0.326103812 | 0.56482837 |
| Figure-4 | Ammonium sulfate concentration | Glucose conc at 0 hours (g/L) | Std. Dev. (0 hours) | Glucose conc at 120 hours (g/L) | Std. Dev. (120 hours) |
| | 5 g/L | 59.90982811 | 0.385083894 | 20.38051569 | 0.975638949 |
| | 8 g/L | 55.14064464 | 1.674869489 | 19.99080566 | 1.047989624 |
| | 15 g/L | 58.30607284 | 1.097949414 | 19.74371196 | 2.30685889 |
| | 30 g/L | 58.86591231 | 1.091677473 | 7.041251396 | 1.717917334 |

**Table-4: Primers used in the study**

| | |
|---|---|
| ylHMGS-Fwd | ccgaccagcactttttgcagtactaaccgcagtcgcaaccccagaacgttggaatc |
| ylHMGS-Rvs | cgtggggacaggccatggaactagtcggtaccctactgcttgatctcgtactttc |
| ylt495HMG1-Fwd | cactttttgcagtactaaccgcagctacgagaagttgtgcgaaccca |
| ylt495HMG1-Rvs | aggccatggaactagtcggtaccctatgaccgtatgcaaatattc |
| ylIDI-F | ccgaccagcactttttgcagtactaaccgcagacgacgtcttacagcgacaaaatc |
| ylIDI-R | cgtggggacaggccatggaactagtcggtaccctacttgatccaccgccgaatctc |
| ylMVK-Fwd | gcactttttgcagtactaaccgcaggactacatcatttcggcgcca |
| ylMVK-Rvs | acaggccatggaactagtcggtaccctaatgggtccagggaccgat |
| ylPMK-Fwd | gcactttttgcagtactaaccgcagaccacctattcggctccggga |
| ylPMK-Rvs | acaggccatggaactagtcggtaccctacttgaacccccttctcgag |
| ylMVD1-Fwd | gcactttttgcagtactaaccgcagatccaccaggcctccaccacc |
| ylMVD1-Rvs | acaggccatggaactagtcggtaccctacttgctgttcttcagaga |
| CrGES-Opt-Fwd | gcactttttgcagtacTAACCGCAGGCTGCTACTATTTCTAACCTG |
| CrGES-Opt-Rvs | aggccatggaactagtcggtaccTTAGAAACAGGGGGTGAAAAAC |
| t3CrGES-Fwd | ttttgcagtacTAACCGCAGTCCTCCTCTTCCTCTTCCTCTTCCATGTCCCTGCCCCTGG |
| CrGPPS-Fwd | actttttgcagtactaaccgcagGCCAAGGCTATCTCCTCTTTCA |
| CrGPPS-Rvs | aggccatggaactagtcggtaccTTACACAGAACACAGGTCGGTCT |
| VoGES-Fwd | actttttgcagtactaaccgcagATCACCTCCTCTTCCTCTGTGC |
| VoGES-Rvs | caggccatggaactagtcggtaccTTAGACGGACACAGAGACGGGG |
| tVoGES-Fwd | cactttttgcagtactaaccgcagTCCTCTCTGCCCGTCTCTAAGT |
| tCrGES_start_FP | TCCTCCTCTTCCTCTTCCTCTTCCATG |
| tCrGES_end_RP2 | CAGAGCCTTGACGTAGTTGTCCACAG |
| EcAtoB_start_seq_FP | AAAAATTGTGTCATCGTCAGTGCGGTAC |
| EcAtoB_end_seq_RP | TTAATTCAACCGTTCAATCACCATCGC |

**Sequence of full length CrGES (*Catharanthus roseus* geraniol synthase) codon optimized for *Yarrowia lipolytica* (1767 bp). Highlighted in green (129 bp) is the chloroplast signal peptide:**

GCTGCTACTATTTCTAACCTGTCTTTTCTGGCTAAGTCCCGAGCTCTGTCCCGACCTTCCTCTTCGTCCCTGTCCTGG
CTGGAGCGACCCAAGACCTCTTCCACCATTTGTATGTCCATGCCCTCTTCTTCCTCCTCTTCCTCTTCCTCTTCCATGT
CCCTGCCCCTGGCCACCCCTCTGATCAAGGACAACGAGTCTCTGATTAAGTTCCTGCGACAGCCCCTGGTGCTGCCT
CACGAGGTGGACGACTCTACCAAGCGACGAGAGCTGCTGGAGCGAACCCGAAAGGAGCTGGAGCTGAACGCCG
AGAAGCCCCTGGAGGCTCTGAAGATGATTGACATCATTCAGCGACTGGGTCTGTCCTACCACTTCGAGGACGACA
TCAACTCCATTCTGACCGGCTTCTCTAACATCTCTTCCCAGACCCACGAGGACCTGCTGACCGCCTCTCTGTGCTTCC
GACTGCTGCGACACAACGGCCACAAGATCAACCCCGACATTTTCCAGAAGTTCATGGACAACAACGGCAAGTTCA
AGGACTCCCTGAAGGACGACACCCTGGGAATGCTGTCCCTGTACGAGGCCTCTTACCTGGGTGCTAACGGCGAGG
AGATCCTGATGGAGGCCCAGGAGTTCACCAAGACCCACCTGAAGAACTCTCTGCCTGCTATGGCTCCTTCCCTGTC
TAAGAAGGTGTCTCAGGCTCTGGAGCAGCCCCGACACCGACGAATGCTGCGACTGGAGGCCCGACGATTCATTGA
GGAGTACGGCGCTGAGAACGACCACAACCCCGACCTGCTGGAGCTGGCCAAGCTGGACTACAACAAGGTCCAGT
CCCTGCACCAGATGGAGCTGTCTGAGATCACCCGATGGTGGAAGCAGCTGGGACTGGTGGACAAGCTGACCTTCG
CTCGAGATCGACCCCTGGAGTGTTTCCTGTGGACCGTGGGTCTGCTGCCTGAGCCTAAGTACTCTGGTTGCCGAAT
TGAGCTGGCCAAGACCATCGCTATTCTGCTGGTCATCGACGACATTTTCGACACCCACGGAACCCTGGACGAGCTG
CTGCTGTTCACCAACGCCATCAAGCGATGGGACCTGGAGGCTATGGAGGACCTGCCCGAGTACATGCGAATCTGT
TACATGGCCCTGTACAACACCACCAACGAGATTTGCTACAAGGTGCTGAAGGAGAACGGATGGTCTGTCCTGCCC
TACCTGAAGGCCACCTGGATCGACATGATTGAGGGTTTCATGGTGGAGGCTGAGTGGTTCAACTCCGACTACGTC
CCCAACATGGAGGAGTACGTGGAGAACGGCGTCCGAACCGCCGGATCTTACATGGCTCTGGTGCACCTGTTCTTC
CTGATCGGCCAGGGAGTGACCGAGGACAACGTCAAGCTGCTGATTAAGCCCTACCCCAAGCTGTTCTCTTCCTCTG
GTCGAATCCTGCGACTGTGGGACGACCTGGGAACCGCCAAGGAAGAGCAGGAGCGAGGTGACCTGGCTTCCTCT
ATCCAGCTGTTCATGCGAGAGAAGGAGATTAAGTCCGAGGAAGAGGGCCGAAAGGGTATTCTGGAGATCATTGA
GAACCTGTGGAAGGAGCTGAACGGAGAGCTGGTCTACCGAGAGGAGATGCCCCTGGCCATCATTAAGACCGCTT
TCAACATGGCCCGAGCTTCTCAGGTGGTCTACCAGCACGAGGAGGATACTTACTTTTCTTCTGTGGACAACTACGT
CAAGGCTCTGTTTTTCACCCCCTGTTTCTAA

**Sequence of full length VoGES (*Valeriana officinalis* geraniol synthase) codon optimized for *Yarrowia lipolytica* (1779 bp). Highlighted in green (138 bp) is the chloroplast signal peptide:**

ATCACCTCCTCTTCCTCTGTGCGATCCCTGTGCTGTCCCAAGACCTCTATCATTTCCGGAAAGCTGCTCCCCTCTCTG
CTCCTGACCAACGTGATCAACGTCTCCAACGGCACCTCCTCCCGAGCCTGCGTGTCTATGTCCTCTCTGCCCGTCTCT
AAGTCCACCGCTTCCTCTATTGCCGCTCCCCTCGTCCGAGACAACGGTTCTGCCCTGAACTTCTTCCCCCAGGCTCCC
CAGGTGGAGATCGACGAGTCCTCTCGAATTATGGAGCTGGTGGAGGCTACCCGACGAACCCTGCGAAACGAGTC
CTCTGACTCTACCGAGAAGATGCGACTCATCGACTCCCTGCAGCGACTGGGACTCAACCACCACTTCGAGCAGGAC
ATTAAGGAGATGCTGCAGGACTTCGCCAACGAGCACAAGAACACCAACCAGGACCTCTTCACCACCTCCCTGCGAT
TCCGACTCCTGCGACACAACGGCTTCAACGTGACCCCCGACGTCTTCAACAAGTTCACCGAGGAGAACGGCAAGTT
CAAGGAGTCCCTGGGAGAGGACACCATCGGCATTCTGTCTCTCTACGAGGCCTCCTACCTGGGCGGCAAGGGAGA
GGAGATCCTGTCTGAGGCTATGAAGTTCTCTGAGTCCAAGCTGCGAGAGTCCTCTGGACACGTCGCTCCCCACATC
CGACGACAGATTTTCCAGTCCCTGGAGCTCCCCCGACACCTCCGAATGGCTCGACTGGAGTCTCGACGATACATCG
AGGAAGACTACTCCAACGAGATTGGCGCCGACTCCTCTCTCCTGGAGCTGGCTAAGCTCGACTTCAACTCTGTGCA
GGCCCTCCACCAGATGGAGCTGACCGAGATTTCTCGATGGTGGAAGCAGCTGGGTCTCTCCGACAAGCTGCCCTT
CGCCCGAGACCGACCCCTGGAGTGCTTCCTGTGGACCGTGGGACTCCTGCCTGAGCCTAAGTACTCCGGTTGTCGA
ATCGAGCTGGCCAAGACCATTGCTGTGCTCCTGGTCATCGACGACATTTTCGACACCTACGGATCTTACGACCAGC
TGATCCTCTTCACCAACGCCATTCGACGATGGGACCTCGACGCTATGGACGAGCTGCCCGAGTACATGAAGATCTG

CTACATGGCCCTCTACAACACCACCAACGAGATTTGTTACAAGGTGCTGAAGGAGAACGGATGGTCTGTCCTGCCT
TACCTGGAGCGAACCTGGATCGACATGGTGGAGGGTTTCATGCTGGAGGCCAAGTGGCTGAACTCCGGAGAGCA
GCCCAACCTGGAGGCTTACATCGAGAACGGTGTGACCACCGCCGGATCTTACATGGCTCTGGTCCACCTCTTCTTC
CTGATTGGAGACGGCGTGAACGACGAGAACGTCAAGCTCCTGCTCGACCCCTACCCCAAGCTCTTCTCCTCTGCCG
GCCGAATCCTGCGACTCTGGGACGACCTGGGAACCGCTAAGGAAGAGCAGGAGCGAGGCGACGTCTCCTCTTCC
ATTCAGCTCTACATGAAGGAGAAGAACGTGCGATCTGAGTCCGAGGGTCGAGAGGGAATCGTCGAGATCATCTA
CAACCTGTGGAAGGACATGAACGGCGAGCTCATTGGTTCCAACGCCCTGCCCCAGGCTATCATTGAGACCTCTTTC
AACATGGCCCGAACCTCCCAGGTCGTGTACCAGCACGAGGACGACACCTACTTCTCTTCCGTGGACAACTACGTCC
AGTCTCTGTTCTTCACCCCCGTCTCTGTGTCCGTC

**Sequence of full length EcAtoB (*E. coli* Acetyl-CoA acetyltransferase) codon optimized for *Yarrowia lipolytica* (1182 bp):**

AAAAATTGTGTCATCGTCAGTGCGGTACGTACTGCTATCGGTAGTTTTAACGGTTCACTCGCTTCCACCAGCGCCAT
CGACCTGGGGGCGACAGTAATTAAAGCCGCCATTGAACGTGCAAAAATCGATTCACAACACGTTGATGAAGTGAT
TATGGGTAACGTGTTACAAGCCGGGCTGGGGCAAAATCCGGCGCGTCAGGCACTGTTAAAAAGCGGGCTGGCAG
AAACGGTGTGCGGATTCACGGTCAATAAAGTATGTGGTTCGGGTCTTAAAAGTGTGGCGCTTGCCGCCCAGGCCA
TTCAGGCAGGTCAGGCGCAGAGCATTGTGGCGGGGGGTATGGAAAATATGAGTTTAGCCCCCTACTTACTCGATG
CAAAAGCACGCTCTGGTTATCGTCTTGGAGACGGACAGGTTTATGACGTAATCCTGCGCGATGGCCTGATGTGCG
CCACCCATGGTTATCATATGGGGATTACCGCCGAAAACGTGGCTAAAGAGTACGGAATTACCCGTGAAATGCAGG
ATGAACTGGCGCTACATTCACAGCGTAAAGCGGCAGCCGCAATTGAGTCCGGTGCTTTTACAGCCGAAATCGTCCC
GGTAAATGTTGTCACTCGAAAGAAAACCTTCGTCTTCAGTCAAGACGAATTCCCGAAAGCGAATTCAACGGCTGAA
GCGTTAGGTGCATTGCGCCCGGCCTTCGATAAAGCAGGAACAGTCACCGCTGGGAACGCGTCTGGTATTAACGAC
GGTGCTGCCGCTCTGGTGATTATGGAAGAATCTGCGGCGCTGGCAGCAGGCCTTACCCCCCTGGCTCGCATTAAA
AGTTATGCCAGCGGTGGCGTGCCCCCCGCATTGATGGGTATGGGGCCAGTACCTGCCACGCAAAAAGCGTTACAA
CTGGCGGGGCTGCAACTGGCGGATATTGATCTCATTGAGGCTAATGAAGCATTTGCTGCACAGTTCCTTGCCGTTG
GGAAAAACCTGGGCTTTGATTCTGAGAAAGTGAATGTCAACGGCGGGGCCATCGCGCTCGGGCATCCTATCGGTG
CCAGTGGTGCTCGTATTCTGGTCACACTATTACATGCCATGCAGGCACGCGATAAAACGCTGGGGCTGGCAACACT
GTGCATTGGCGGCGGTCAGGGAATTGCGATGGTGATTGAACGGTTGAATTAA

**Sequence of full length HMGS (HMG synthase -1338 bp):**

tcgcaaccccagaacgttggaatcaaagccctcgagatctacgtgccttctcgaattgtcaaccaggctgagctcgagaagcacgacggtgtcgctgc
tggcaagtacaccattggtcttggtcagaccaacatggcctttgtcgacgacagagaggacatctattcctttgccctgaccgccgtctctcgactgctc
aagaacaacaacatcgaccctgcatctattggtcgaatcgaggttggtactgaaacccttctggacaagtccaagtccgtcaagtctgtgctcatgcag
ctctttggcgagaacagcaacattgagggtgtggacaacgtcaacgcctgctacggaggaaccaacgccctgttcaacgctatcaactgggttgagg
gtcgatcttgggacggccgaaacgccatcgtcgttgccggtgacattgccctctacgcaaagggcgctgcccgacccaccggaggtgccggctgtgtt
gccatgctcattggccccgacgctcccctggttcttgacaacgtccacggatcttacttcgagcatgcctacgatttctacaagcctgatctgacctccga
gtaccccctatgttgatggccactactccctgacctgttacacaaaggccctcgacaaggcctacgctgcctacaacgcccgagccgagaaggtcggtct
gttcaaggactccgacaagaagggtgctgaccgatttgactactctgccttccacgtgcccacctgcaagcttgtcaccaagtcttacgctcgacttctc
tacaacgactacctcaacgacaagagcctgtacgagggccaggtccccgaggaggttgctgccgtctcctacgatgcctctctcaccgacaagaccgt
cgagaagaccttccttggtattgccaaggctcagtccgccgagcgaatggctccttctctccagggacccaccaacaccggtaacatgtacaccgcctc
tgtgtacgcttctctcatctctctgctgacttttgtccccgctgagcagctgcagggcaagcgaatctctctcttctcttacggatctggtcttgcttccact
cttttctctctgaccgtcaagggagacatttctcccatcgtcaaggcctgcgacttcaaggctaagctcgatgaccgatccaccgagactcccgtcgact
acgaggctgccaccgatctccgagagaaggcccacctcaagaagaactttgagccccagggagacatcaagcacatcaagtctggcgtctactacct
caccaacatcgatgacatgttccgacgaaagtacgagatcaagcagtag

**Sequence of t495HMG1 (HMG reductase- 1518 bp):**

ctacgagaagttgtgcgaacccagtctgtgaaggtggttgagaagcacgttcctatcgtcattgagaagcccagcgagaaggaggaggacacctctt
ctgaagactccattgagctgactgtcggaaagcagcccaagcccgtgaccgagacccgttctctggacgacctagaggctatcatgaaggcaggtaa
gaccaagcttctggaggaccacgaggttgtcaagctctctctcgagggcaagcttcctttgtatgctcttgagaagcagcttggtgacaacacccgagc
tgttggcatccgacgatctatcatctcccagcagtctaataccaagactttagagacctcaaagcttccttacctgcactacgactacgaccgtgtttttg
gagcctgttgcgagaacgttattggttacatgcctctccccgttggtgttgctggccccatgaacattgatggcaagaactaccacattcctatggccac
cactgagggttgtcttgttgcctcaaccatgcgaggttgcaaggccatcaacgccggtggcggtgttaccactgtgcttactcaggacggtatgacacg
aggtccttgtgtttccttcccctctctcaagcgggctggagccgctaagatctggcttgattccgaggagggtctcaagtccatgcgaaaggccttcaac
tccacctctcgatttgctcgtctccagtctcttcactctacccttgctggtaacctgctgtttattcgattccgaaccaccactggtgatgccatgggcatga
acatgatctccaagggcgtcgaacactctctggccgtcatggtcaaggagtacggcttccctgatatggacattgtgtctgtctcgggtaactactgcac
tgacaagaagcccgcagcgatcaactggatcgaaggccgaggcaagagtgttgttgccgaagccaccatccctgctcacattgtcaagtctgttctca
aaagtgaggttgacgctcttgttgagctcaacatcagcaagaatctgatcggtagtgccatggctggctctgtgggaggtttcaatgcacacgccgcaa
acctggtgaccgccatctaccttgccactggccaggatcctgctcagaatgtcgagtcttccaactgcatcacgctgatgagcaacgtcgacggtaacc
tgctcatctccgtttccatgccttctatcgaggtcggtaccattggtggaggtactattttggagccccagggggctatgctggagatgcttggcgtgcga
ggtcctcacatcgagacccccggtgccaacgcccaacagcttgctcgcatcattgcttctggagttcttgcagcggagctttcgctgtgttctgctcttgc
tgccggccatcttgtgcaaagtcatatgacccacaaccggtcccaggctcctactccggccaagcagtctcaggccgatctgcagcgtctacaaaacg
gttcgaatatttgcatacggtcatag

**Sequence of IDI1 (IPP isomerase - 810 bp):**

acgacgtcttacagcgacaaaatcaagagtatcagcgtgagctctgtggctcagcagtttcctgaggtggcgccgattgcggacgtgtccaaggctag
ccggcccagcacggagtcgtcggactcgtcggccaagctatttgatggccacgacgaggagcagatcaagctgatggacgagatctgtgtggtgctg
gactgggacgacaagccgattggcggcgcgtccaaaaagtgctgtcatctgatggacaacatcaacgacggactggtgcatcgggccttttccgtgtt
catgttcaacgaccgcggtgagctgcttctgcagcagcgggcggcggaaaaaatcacctttgccaacatgtggaccaacacgtgctgctcgcatcctc
tggcggtgcccagcgagatgggcgggctggatctggagtcccggatccagggcgccaaaaacgccgcggtccggaagcttgagcacgagctgggaa
tcgaccccaaggccgttccggcagacaagttccatttcctcacccggatccactacgccgcgccctcctcgggcccctggggcgagcacgagattgact
acattctgtttgtccggggcgaccccgagctcaaggtggtggccaacgaggtccgcgataccgtgtgggtgtcgcagcagggactcaaggacatgatg
gccgatcccaagctggttttcaccccttggttccggctcatttgtgagcaggcgctgtttccctggtgggaccagttggacaatctgcccgcgggcgatg
acgagattcggcggtggatcaagtag

**Sequence of ERG12 (Mevalonate kinase - 1347 bp):**

gactacatcatttcggcgccaggcaaagtgattctatttggtgaacatgccgctgtgtttggtaagcctgcgattgcagcagccatcgacttgcgaacat
acctgcttgtcgaaaccacaacatccgacaccccgacagtcacgttggagtttccagacatccacttgaacttcaaggtccaggtggacaagctggca
tctctcacagcccagaccaaggccgaccatctcaattggtcgactcccaaaactctggataagcacattttcgacagcttgtctagcttggcgcttctgg
aagaacctgggctcactaaggtccagcaggccgctgttgtgtcgttcttgtacctctacatccacctatgtccccccttctgtgtgcgaagattcatcaaac
tgggtagttcgatcaacgctgcctatcggcgcgggcctgggctcttccgcatccatttgtgtctgtttggctgcaggtcttctggttctcaacggccagctg
agcattgaccaggcaagagatttcaagtccctgaccgagaagcagctgtctctggtggacgactggtccttcgtcggtgaaatgtgcattcacggcaa
cccgtcgggcatcgacaatgctgtggctactcagggaggtgctctgttgttccagcgacctaacaaccgagtccctcttgttgacattcccgagatgaa
gctgctgcttaccaatacgaagcatcctcgatctaccgcagacctggttggtggagtcggagttctcactaaagagtttggctccatcatggatcccatc
atgacttcagtaggcgagatttccaaccaggccatggagatcatttctagaggcaagaagatggtggaccagtctaaccttgagattgagcagggtat
cttgcctcaacccacctctgaggatgcctgcaacgtgatggaagatggagctactcttcaaaagttgagagatatcggttcggaaatgcagcatctagt
gagaatcaatcacggcctgcttatcgctatgggtgtttcccacccgaagctcgaaatcattcgaactgcctccattgtccacaacctgggtgagaccaa
gctcactggtgctggaggaggaggttgcgccatcactctagtcacttctaaagacaagactgcgacccagctggaggaaaatgtcattgctttcacag
aggagatggctacccatggcttcgaggtgcacgagactactattggtgccagaggagttggtatgtgcattgaccatccctctctcaagactgttgaag
ccttcaagaaggtggagcgggcggatctcaaaaacatcggtccctggacccattag

**Sequence of ERG8 (Phosphomevalonate kinase - 1254 bp):**

accacctattcggctccgggaaaggccctcctttgcggcggttatttggttattgatccggcgtattcagcatacgtcgtgggcctctcggcgcgtattta
cgcgacagtttcggcttccgaggcctccaccacctctgtccatgtcgtctctccgcagtttgacaagggtgaatggacctacaactacacgaacggcca
gctgacggccatcggacacaacccatttgctcacgcggccgtcaacaccgttctgcattacgttcctcctcgaaacctccacatcaacatcagcatcaa
aagtgacaacgcgtaccactcgcaaattgacagcacgcagagaggccagtttgcataccacaaaaaggcgatccacgaggtgcctaaaacgggcct
cggtagctccgctgctcttaccaccgttcttgtggcagctttgctcaagtcatacggcattgatcccttgcataacacccacctcgttcacaacctgtccc
aggttgcacactgctcggcacagaagaagattgggtctggatttgacgtggcttcggccgtttgtggctctctagtctatagacgtttcccggcggagtc
cgtgaacatggtcattgcagctgaagggacctccgaatacggggctctgttgagaactaccgttaatcaaaagtggaaggtgactctggaaccatcct
tcttgccgccgggaatcagcctgcttatgggagacgtccagggaggatctgagactccaggtatggtggccaaggtgatggcatggcgaaaagcaaa
gccccgagaagccgagatggtgtggagagatctcaacgctgccaacatgctcatggtcaagttgttcaacgacctgcgcaagctctctctcactaaca
acgaggcctacgaacaacttttggccgaggctgctcctctcaacgctctaaagatgataatgttgcagaaccctctcggagaactagcacgatgcatt
atcactattcgaaagcatctcaagaagatgacacgggagactggtgctgctattgagccggatgagcagtctgcattgctcaacaagtgcaacactta
tagtggagtcattggaggtgttgtgcctggagcaggaggctacgatgctatttctcttctggtgatcagctctacggtgaacaatgtcaagcgagagag
ccagggagtccaatggatggagctcaaggaggagaacgagggtctgcggctcgagaaggggttcaagtag

**Materials:**

| Material | Company |
|---|---|
| *E. coli* Dh5α | NEB |
| *E. coli* Dh10β | NEB |
| CSM/ CSM-Leu/ CSM-Leu-Ura | Sunrise science products |
| Yeast nitrogen base without amino acids | BD Difco |
| Gibson assembly mastermix | NEB |
| Restriction enzymes | NEB |
| ZR plasmid mini-prep kit | Zymo research (D4054) |
| PCR cleanup kit | Zymo research |
| gDNA extraction kit | Thermo Scientific (PI78870) |
| Q5 High-Fidelity 2X Master Mix for PCR | NEB (M0492S) |
| Qscript™ One-Step SYBR® Green qRT-PCR Kit, QuantaBio for qPCR | VWR (95054-946) |
